# A Novel Model for Proton Transport Mediated by Uncoupling Protein 1

**DOI:** 10.1101/2025.04.28.650983

**Authors:** Luise Jacobsen, Sneha Menon, Michael J Gaudry, Ali Asghar Hakami Zanjani, Peter Reinholdt, Martin Jastroch, Himanshu Khandelia

## Abstract

Uncoupling Protein 1 (UCP1) is a mitochondrial protein which drives thermogenesis in brown adipose tissue. UCP1 facilitates the dissipation of the proton gradient as heat and plays a critical role in energy expenditure and metabolic regulation. We employ advanced molecular simulations and mutagenesis to reveal the mechanism of UCP1-mediated proton and fatty acid (FA) transport. We demonstrate that FAs bind spontaneously to UCP1’s central substrate-binding site. In the binding site, a proton transfer to the FA is facilitated by a key aspartate residue (D28) and a coordinating water molecule. The protonated FA exits UCP1 through a well defined pathway, and releases its proton into the mitochondrial matrix. UCP1 then facilitates the return of deprotonated FAs to the intermembrane space. Nucleotide binding disrupts this mechanism by inducing conformational changes in the transmembrane helices and obstructing the FA return pathway. Our mechanism explains every step of the transport cycle, is supported by simulation and biochemical data, and explains a diverse set of biochemical data about the transport mechanisms in UCP1 and its analogues: ANT, UCP2 and UCP3.

## Introduction

The prevalence of obesity has more than doubled since 1990, with one in eight individuals affected by obesity in 2022, according to the World Health Organization [WHO, 2024]. Obesity is associated with several major health conditions, including cardiovascular disease, diabetes, and cancer [Bray et al., 2017], contributing to over 5 million deaths annually [GBD 2019 Risk Factors Collaborators, 2020]. This public health crisis imposes a significant economic burden, costing an average of 1.8% of the gross domestic product across eight geographically and economically diverse countries[Okunogbe et al., 2021]. Obesity is now considered a pandemic [The Lancet Gastroenterology Hepatology, 2021].

Brown adipose tissue (BAT) and beige adipose tissue are promising therapeutic targets for obesity treatment. Unlike white adipose tissue, which stores energy as fat, BAT actively clears blood glucose [Cannon and Nedergaard, 2004, Stanford et al., 2013, Keipert and Jastroch, 2014] and triggers non-shivering thermogenesis, a process fueled by the metabolic rate and energy expenditure. Therefore, increased BAT activation has potential to improve metabolic health and combat obesity and obesity-related diseases.

The thermogenic function of BAT is attributed to a high concentration of the uncoupling protein 1 (UCP1), an inner mitochondrial membrane (IMM) protein [Cannon and Nedergaard, 2004, Shabalina et al., 2004]. UCP1 facilitates proton leakage from the intermembrane space (IMS) to the mitochondrial matrix without ATP synthesis, directly dissipating the electrochemical gradient across the IMM as heat. The uncoupling from ATP homeostasis enables the acceleration of oxidation, and thereby, heat output. Targeting UCP1 for therapeutic activation is thus a promising avenue for obesity treatment.

Fatty acids (FAs) physiologically activate UCP1 during adrenergic stimulated lipolysis, such as during acute cold exposure [Cannon and Nedergaard, 2004]. Other known mitochondrial uncouplers, like 2,4-Dinitrophenol (DNP), have also been proposed to activate UCP1 via a mechanism similar to that of FAs. However, the involvement of UCP1 in DNP induced mitochondrial uncoupling is still debated [Bertholet et al., 2022, Shabalina et al., 2025]. On the other hand, it is widely accepted that the uncoupling activity of UCP1 is inhibited by purine nucleotides (ATP, ADP, GTP, GDP) [Keipert and Jastroch, 2014, Shabalina et al., 2004].

Cryogenic electron microscopy (cryo-EM) structures of apo, DNP-bound, ATP-bound, and GTP-bound UCP1 have been resolved [Kang and Chen, 2023, Jones et al., 2023] and mutagenesis has identified residues involved in proton transport [Bienengraeber et al., 1998, Klingenberg et al., 1999, Urbánková et al., 2003a, Jiménez-Jiménez et al., 2006, Echtay et al., 2000, Gagelin et al., 2023]. However, the molecular mechanisms underlying UCP1 activation by FAs and UCP1 inhibition by nucleotides remain unresolved. Importantly, although FA is the most relevant UCP1 ligand, no FA-bound UCP1 structure is yet resolved. An FA-bound state is expected to provide insights into the UCP1 active (proton-conducting) state. The difficulty in trapping and resolving an FA-bound state suggests that FA may not bind UCP1 tightly but instead activates UCP1 through an alternative mechanism.

Like all other members of the solute carrier 25 (SLC25) protein family, UCP1 is ∼300 residues in length and exhibits a threefold symmetric structure: six transmembrane helices (TM1-TM6) fold in to three pairs connected by loops (L1-L3) and short matrix-facing helices (h1-h3, Fig. 1)[Kunji et al., 2020]. SLC25 proteins typically alternate between two major conformations: open toward the matrix (m-state) or open toward the IMS/cytosol (c-state). Intramolecular salt bridge networks unique to each state act as gates to keep the protein conformation open to only one side. However, it is possible that UCP1 never adopts an m-state conformation. First, while residues involved in these salt bridge networks are present in UCP1 [Jones et al., 2024], current experimental structures (apo and ligand-bound) exclusively capture the c-state [Jones et al., 2023, Kang and Chen, 2023]. Secondly, FAs and nucleotides approach UCP1 from the IMS in the c-state [Fedorenko et al., 2012, Gagelin et al., 2023]. Finally, most SLC25 transporters such as the Adenine Nucleotide Translocator (ANT) transport two different substrates across the membrane in opposite directions. Such transport necessitates an alternating-access transport mechanism with two conformational states (c- and m-state). UCP1, on the other hand, only transports protons in one direction, and such transport can in principle be achieved by carefully orchestrated conformational changes in the c-state only.

**Figure 1:**
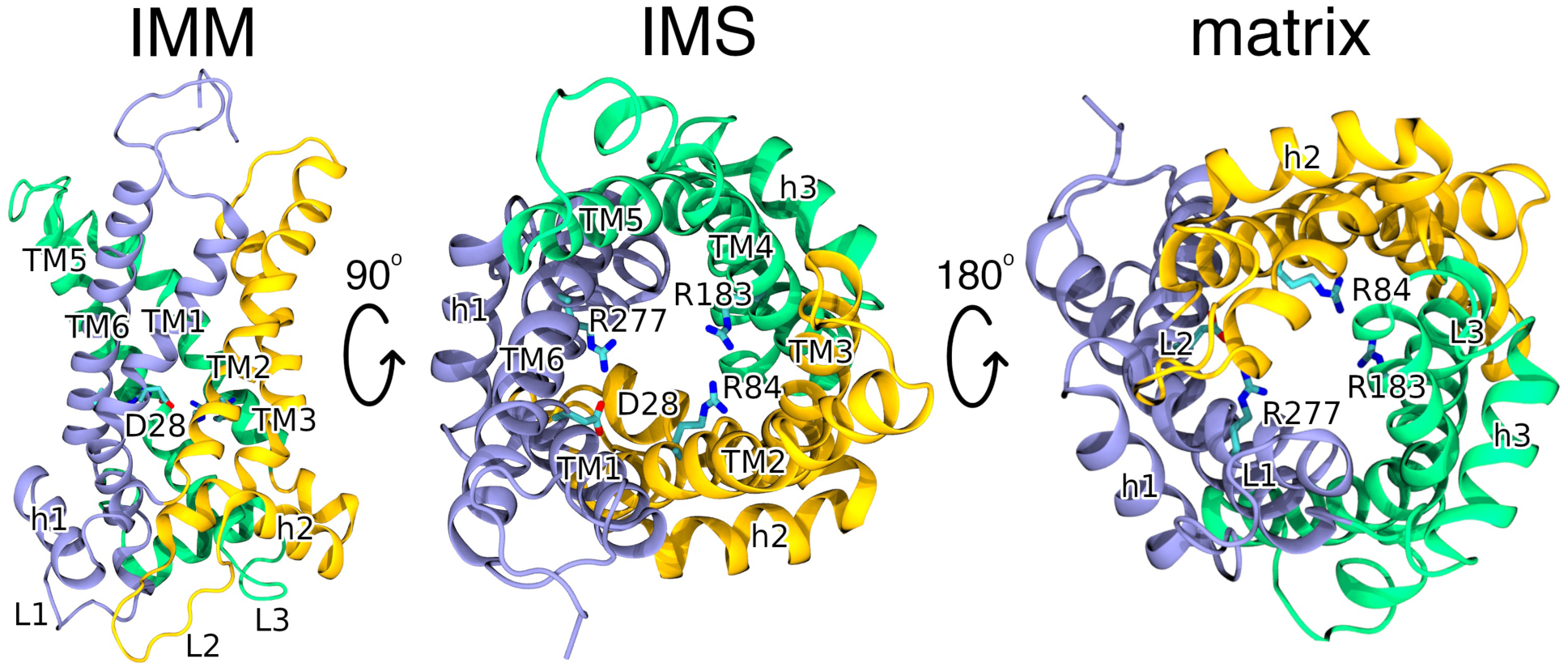
Structural views of uncoupling protein 1 (UCP1) UCP1 is seen from three perspectives: the inner mitochondrial membrane (IMM), the inter-membrane spaces (IMS), and the matrix. UCP1 is comprised of six transmembrane helices (TM1-TM6), organized into three pairs connected by matrix-facing loops (L1-L3) and short helices (h1-h3). Key residues at the central substrate-binding site: R84, R183, R277, and aspartate D28 are shown in stick representation.

Several transport models have been proposed for UCP1-mediated proton transport [Bertholet and Kirichok, 2017]: In the *Buffering Model*, the carboxyl group of FA binds within UCP1 and forms a proton translocation pathway together with titratable residues [Winkler and Klingenberg, 1994, Klingenberg and Huang, 1999]. In the *Shuttling Model*, the carboxyl group resides inside UCP1 while the hydrophobic tail of the FA remains anchored in the membrane, allowing the carboxyl group to shuttle protons between the IMS and matrix [Fedorenko et al., 2012]. In the *Cycling Model*, the FA itself carries protons across the membrane independently of UCP1. FAs protonate in the IMS leaflet, flip to the matrix leaflet, and release the proton. In this model, UCP1 only facilitates the return of the anionic fatty acid to the IMS leaflet [Skulachev, 1991].

Here, we present extensive simulation-based evidence to support a novel proton transport mechanism through UCP1, which does not necessitate a transition to the m-state. We apply three different acronyms for fatty acids: FA for fatty acids in general or when the state is ambiguous as in constant pH simulations, FA*^−^* for deprotonated and anionic FA, and FAp for protonated and neutral FA. Starting from the apo state (Fig. 2A), we first demonstrate that FA*^−^* can spontaneously bind to the central substrate-binding site of UCP1 (residues R84, R183, and R277; Fig. 2B). Next, FA*^−^* gets protonated by a key aspartate residue (D28; Fig. 2C), with the proton transfer mediated through a water molecule bridging FA*^−^* and D28. The protonated FAp then exits UCP1 from between TM5 and TM6 (Fig. 2D), enters the matrix leaflet, and releases its proton to the matrix space (Fig. 2E). Finally, the then anionic FA*^−^* flips back to the IMS leaflet sliding along a polar pathway on TM2, completing the cycle (Fig. 2F). Nucleotides inhibit proton transport by compromising the polar pathway return route.

**Figure 2:**
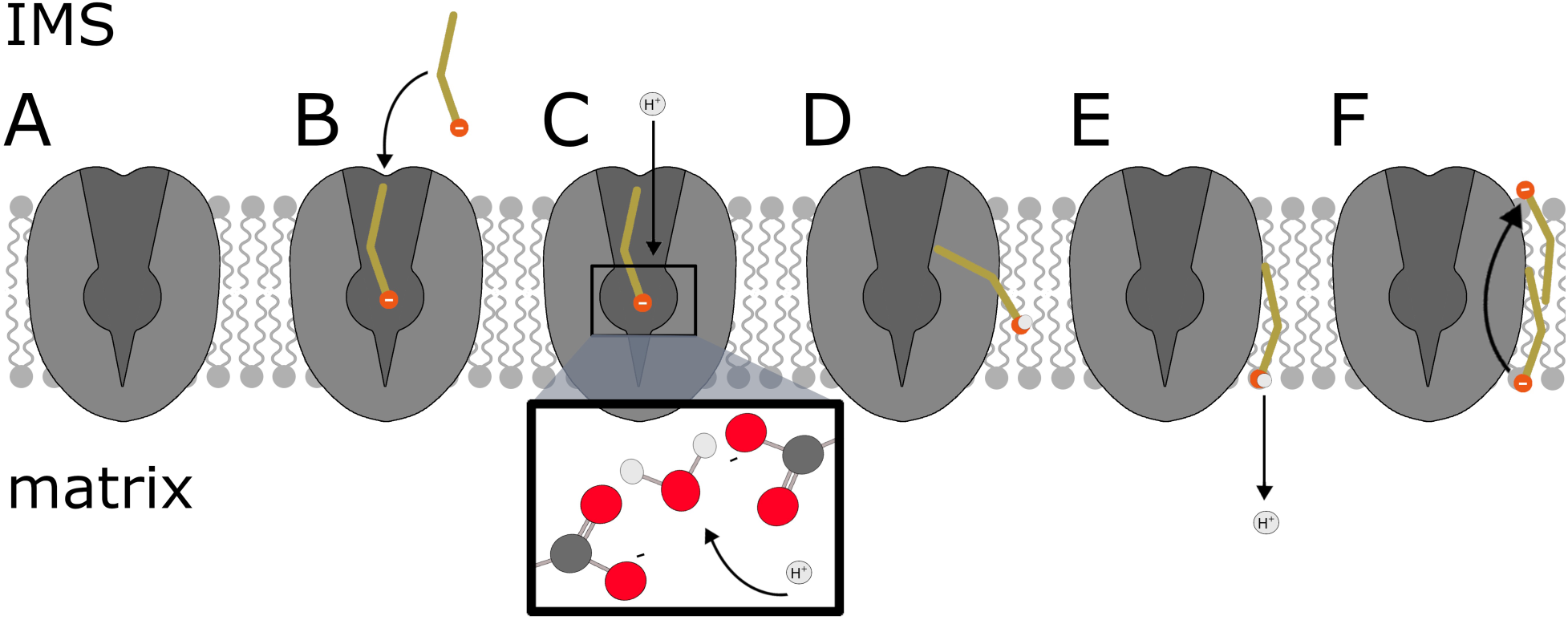
Proposed mechanism for proton and fatty acid (FA) transport by UCP1. **A**: In the apo state, **B**: an anionic FA (FA*^−^*) enters UCP1 and binds to the central substrate-binding site. **C**: A water molecule bridges FA*^−^* and an aspartate residue (D28) in the binding site. The water molecule facilitates protonation of FA*^−^*. **D**: The protonated FA (FAp) exits UCP1 between TM5 and TM6 and enters the matrix leaflet. **E**: Once in the matrix leaflet, FAp releases its proton into the matrix. **F**: To complete the transport cycle, FA*^−^* is transported back to the IMS leaflet via a polar pathway along TM2.

To elucidate the molecular details of the mechanism, we employed a range of simulation techniques: standard all-atom and coarse-grained molecular dynamics (MD), constant pH simulations, free energy perturbation (FEP), umbrella sampling, and quantum mechanics/molecular mechanics (QM/MM). Finally, experimental mutagenesis was performed to validate the roles of specific residues predicted to participate in the mechanism proposed by the simulations. In the following section, we will present evidence for each step in the proposed transport mechanism (Fig. 2A-E).

## Results

### Preliminary Simulations of Apo and ATP-Bound UCP1

All simulations described below were based on well-equilibrated 300 ns preliminary simulations of the cryo-EM structures of UCP1 in the apo (pdb 8hbv, PRE-APO) or ATP-bound (pdb 8hbw, PRE-ATP) states [Kang and Chen, 2023] under physiological conditions (Figs. S1 and S2).

### Substrate Binding Alters the Ionization State of D28

Proton transfer events in proteins occur via protonation and deprotonation of amino acids. The ionization states and pKa values of amino acids within a substrate-binding site can differ significantly from their values in water and may be altered by the binding of charged substrates [Yamamoto et al., 2019, Han et al., 2017, Rui et al., 2016]. To investigate possible pKa shifts of the ionizable residues in UCP1’s substrate-binding site, we launched 1 µs constant pH simulations of UCP1 in the apo (CpH-APO), ATP-bound (CpH-ATP), and FA-bound (CpH-FA) states. Lysine, glutamate, and aspartate residues exposed to the IMS (D8, D28, D35, D97, D196, D210, D211, D234, D304, K38, K107, K116, K138, K199, K204, K237, K293, E101, E108, E135, E191, E200, and E290) were kept titratable. The simulations were conducted at pH 6.8 to mimic the IMS conditions.

Most titratable amino acids retained their default ionization states in water during the simulations. However, the ionization states of D28 and E191 deviated from their values in water (Fig. 3A,B). D28 was consistently deprotonated (D28*^−^*) in CpH-APO, but never in CpH-FA and CpH-ATP. In contrast, D28 got protonated (D28p) in CpH-FA (Fig. 3A). Titration curves derived from constant pH simulations over a pH range from 1 to 9 revealed that D28 remained deprotonated in apo-UCP1 even at pH 1. In contrast, D28 was fully protonated across the entire pH range (1-9) in FA-bound UCP1 (Fig. S3).

**Figure 3:**
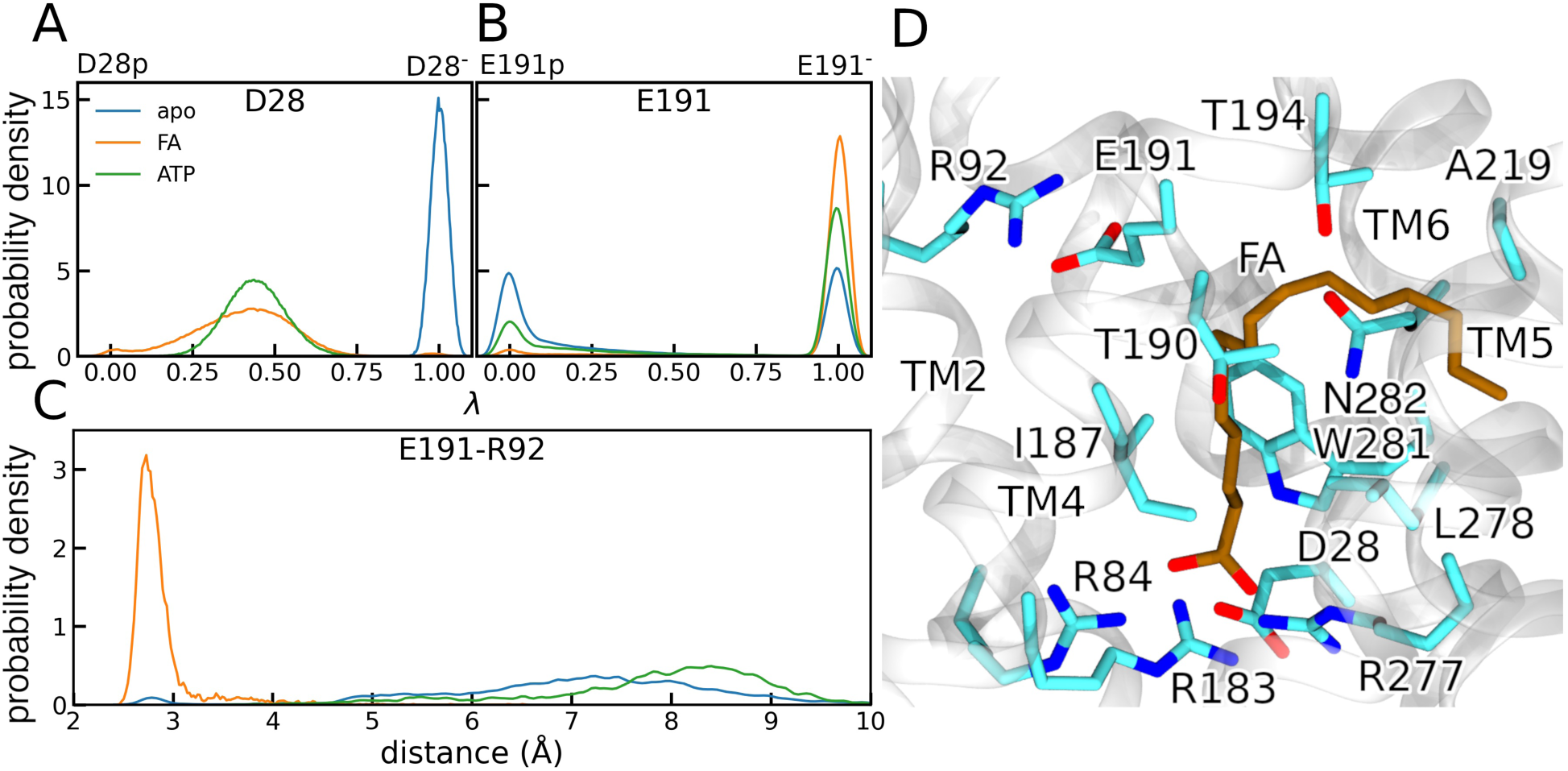
Protonation states and gate formation in constant pH simulations. Probability density of *λ* values for **A**: D28 and **B**: E191, calculated from the final 200 ns of 1 µs constant pH simulations of apo (CpH-APO), FA-bound (CpH-FA), and ATP-bound (CpH-ATP) UCP1. *λ* = 0 indicates a fully protonated residue, while *λ* = 1 indicates complete deprotonation. **C**: Probability density of the distance between the E191 carboxyl group and the R92 amine in simulations CpH-APO, CpH-FA, and CpH-ATP. **D**: Structural snapshot from the CpH-FA simulation, depicting FA*^−^* bound to the arginine triplet (R84, R183, and R277) with the R92-E191 gate closed and the FA*^−^* tail penetrating UCP1 between TM5 and TM6.

E191 had a 50% likelihood of being protonated (E191p) in CpH-APO (Fig. 3B). Although both protonation states were possible in CpH-FA and CpH-ATP, the anionic state (E191*^−^*) was more prevalent, particularly in CpH-FA. In CpH-FA, the predominantly anionic E191 kept the R92-E191 gate closed (Fig. 3C,D). It is worth noting that the R92-E191 gate did not impede the access of FA*^−^* to the binding site (Fig. 3D), in agreement with previous studies which showed that proton transport is independent of R92 and E191 [Echtay et al., 2001, Winkler et al., 1997, Echtay et al., 1997].

The FA remained deprotonated throughout the 1 µs CpH-FA simulation. However, in a longer 5 µs simulation, the FA momentarily got protonated under specific conditions: when D28 was deprotonated, a salt bridge formed between D28*^−^* and R277. The salt bridge allowed the FA to approach D28*^−^* (*<*6 Å) and get protonated (Fig. S4). The FA approached D28 only when the R277-D28*^−^* salt bridge was present.

The FA remained bound to UCP1 throughout the 5 µs CpH-FA simulation, maintaining stable hydrophobic interactions with L278, I187, and W281, which stabilized carbons 1 and 2 of the FA (Fig. 3D). T190, N282, T194, and A219 stabilized the remainder of the FA chain (occupancy*>*50%) which penetrated UCP1 between TM5 and TM6. The carboxyl group of the FA formed stable electrostatic interactions with the arginine triplet (R84, R183, and R277). The following section describes how the interactions between the FA and D28 can facilitate a proton transfer to the FA.

### FA**^−^** Spontaneously Approaches D28**^−^**

Constant pH simulations indicated that D28 was protonated (D28p) in FA-bound UCP1. Based on this, we hypothesized that D28p could directly transfer a proton to the FA. To test this, we launched five conventional 1.5 µs simulations with an anionic FA (FA*^−^*) at the opening of UCP1. We kept D28 protonated in these simulations. In all five replicas, FA*^−^* bound tightly to R84 and R183 but had limited binding to other residues. The carboxyl group of FA*^−^* was at least 8 Å from D28p (Fig. S5), indicating that a direct proton transfer between D28p and FA*^−^* was impossible.

We then speculated that a water molecule (hydronium ion) could act as a bridge between FA*^−^* and D28*^−^* and facilitate a proton transfer. Such proton transfer events through intervening water molecules are documented in the cyanobacterial photosystem II [Kemmler et al., 2019] and bacteriorhodopsin [Luecke et al., 1999]. To explore this possibility, we conducted a 1.1 µs conventional simulation where we positioned the FA*^−^* at the opening of UCP1 and kept D28 deprotonated (D28*^−^*). During the simulation, FA*^−^* spontaneously diffused to the arginine triplet (R84, R183 and R277; Fig. S6). Upon binding, the distance between the carboxylate groups of FA*^−^* and D28*^−^* stabilized at ∼4.9 Å (Fig. S6 and S7), a distance close enough for a water molecule to bridge FA*^−^* and D28*^−^*. We detected such a bridging water molecule in 20% of the simulation time. A bridged water molecule was defined as one positioned within 3 Å of both the FA*^−^* and D28*^−^* carboxylate groups.

Knowing that a water molecule can bridge FA*^−^* and D28*^−^*, we wanted to investigate the possibility of a proton transfer via such a water molecule. We employed QM/MM simulations to study this proton transfer, because such events cannot be accessed by conventional MD simulations. The results of these QM/MM simulations are described in the next section.

### A Water Molecule Facilitates Proton Transfer from D28p to FA^−^

We selected five trajectory frames with water coordinated between FA*^−^* and D28*^−^*, and used these as initial conformations to launch five independent QM/MM simulations (QM1-QM5; Fig. 4A). We initially placed the excess proton on D28. In simulations QM1 and QM4, the excess proton was rapidly transferred to the FA via the water molecule within 1.7 ps (Figs. 4B and S8B, and Supplementary video 1). In the simulation QM3, complete proton transfers occurred briefly (*<*1 ps) on two occasions during the 45 ps simulation (Fig. S8A). No proton transfer was observed within 25 ps in simulations QM2 and QM5. Overall, the QM/MM simulations demonstrated that a proton can get transferred from D28p to the FA through a water molecule.

**Figure 4:**
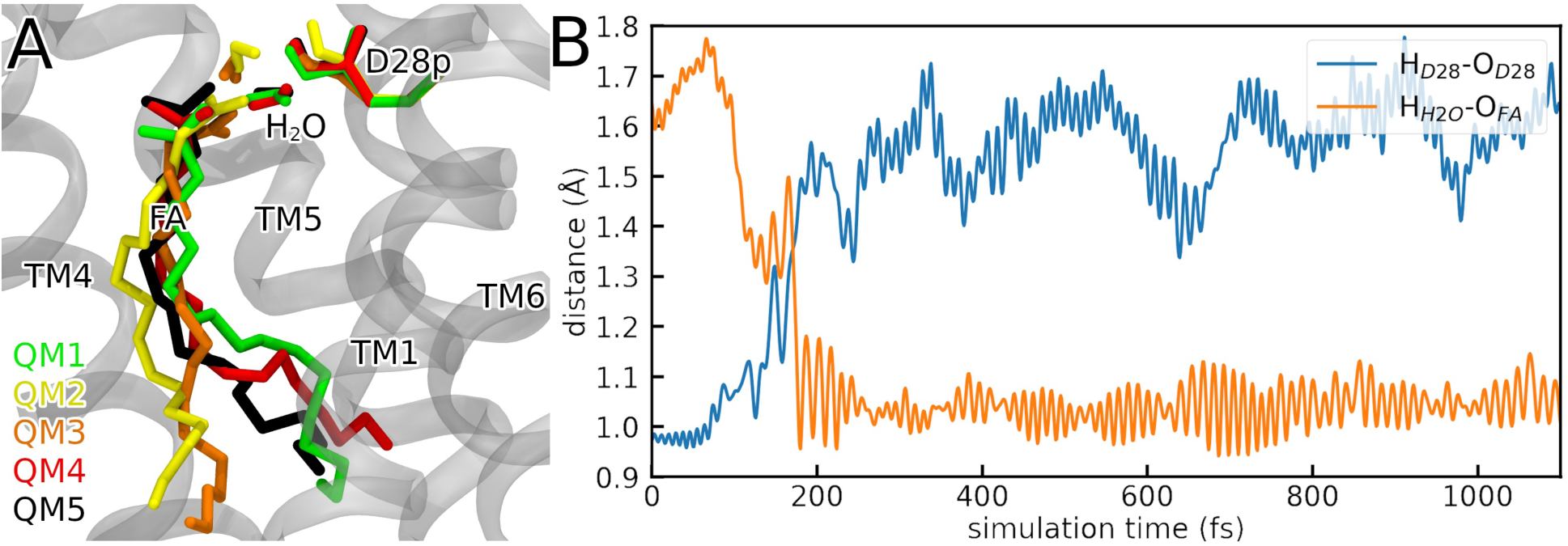
Proton transfer in QM/MM simulations. **A**: Starting conformations for QM/MM simulations. **B**: Distances between the transferring hydrogen atoms (H_D28_, initially on D28, and H_H2O_, initially on the water molecule) and the oxygen atoms of D28 (O_D28_) or FA (O_FA_) in simulation QM1.

To further quantify the thermodynamic feasibility of a proton transfer, we performed free energy calculations using the free energy perturbation (FEP) method. For the starting conformation, we used a simulation frame with a coordinated water molecule between FA*^−^* and D28*^−^*. We placed the proton initially on D28 (D28p) and then moved the proton to the FA during FEP. We maintained coordination of water to both FA and D28 by using distance restraints between the water oxygen atom and the oxygen atoms of the FA and D28. The free energy change associated with a proton transfer from D28 to the FA was −1.3 ± 0.4 kcal/mol indicating that the proton transfer was energetically favourable. While the free energy values for the forward (−0.6 ± 0.5 kcal/mol) and backward (1.6 ± 0.1 kcal/mol) transformations did not completely converge – a common occurrence in FEP calculations [Razavi et al., 2017, Jacobsen et al., 2021], both indicated a favourable transfer of the proton from D28 to the FA. Once protonated, the FA is expected to spontaneously migrate to the matrix-facing leaflet of the membrane (Fig. 2D). In the next set of simulations, we investigated the possibility of FAp exiting the UCP1 binding cavity and approaching the matrix leaflet.

### FAp Spontaneously Diffuses to the Matrix Leaflet

We simulated six different FAp-bound UCP1 structures to investigate FA dynamics after protonation in the binding site. The starting conformations were extracted from previous simulations of FA*^−^*-bound UCP1 with D28 deprotonated (D28*^−^*). Without strong electrostatic interactions with the arginine triplet at the binding site, FAp diffused into the membrane through the gap between TM5 and TM6. The protonated carboxyl acid group of FAp was attracted to G279, S280, and G223 located in the center of the IMM (Fig. 5A,B).

**Figure 5:**
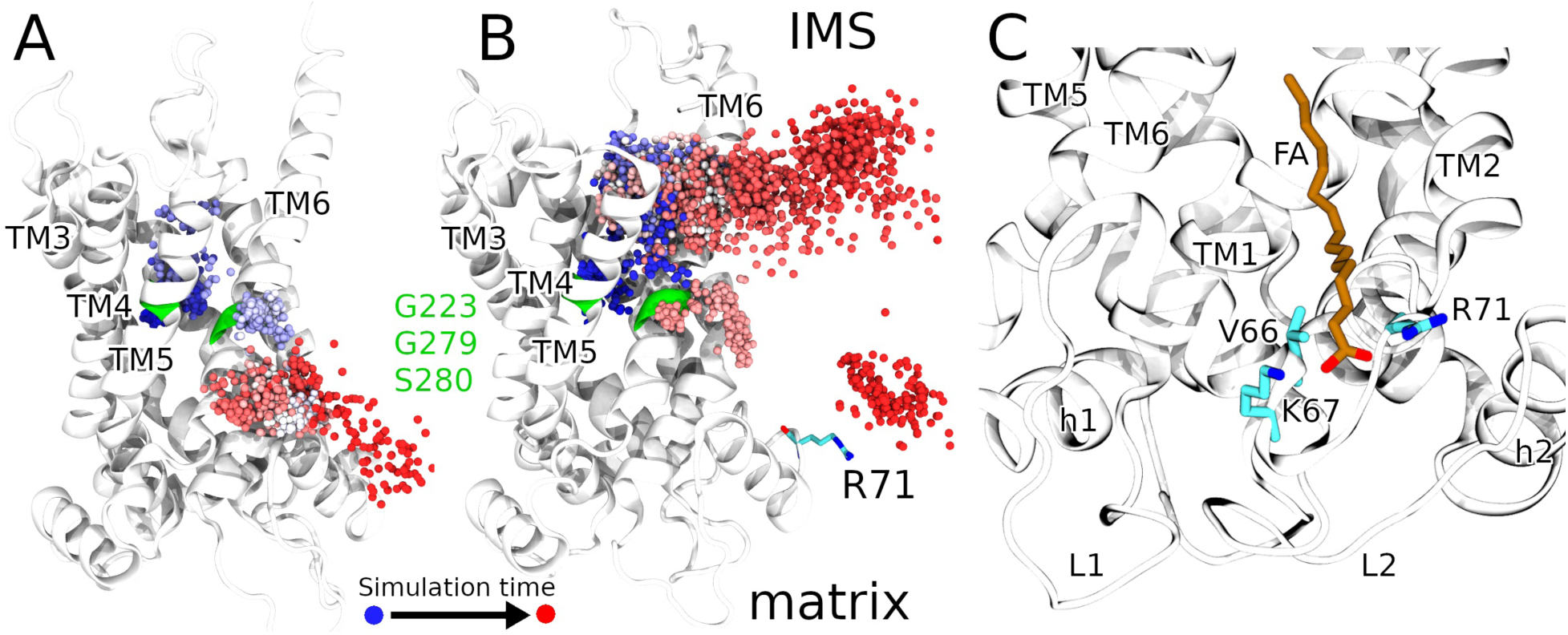
Migration of FAp from the central substrate-binding site to the matrix. **A**,**B**: Position of the carboxylic acid carbon atom of FAp relative to UCP1 across different simulation snapshots. The simulation time is color coded such that the simulation progressed from blue → red spheres. The polar residues G279, S280, and G223 are shown in green cartoon, and R71 is represented in sticks in panel B. Only trajectory frames where FAp was near the protein are included, with 250 ps intervals between two spheres. Corresponding figures for other simulations are available in Fig. S9. **C**: the binding of FA between R71 and K67 on the exterior of UCP1 in a constant pH simulation.

In two of the six simulations, FAp migrated toward the matrix leaflet, where it transiently remained near TM6 before detaching from UCP1 and diffusing into the membrane (Fig. 5A). In three simulations, FAp initially diffused to the IMS leaflet, but rapidly flipped to the matrix leaflet in two of these cases. Within the matrix leaflet, FAp transiently interacted with R71 before diffusing into the membrane (Fig. 5B). In the final simulation, FA*^−^* remained in the central UCP1 binding site throughout the 2.5 µs simulation. Overall, FAp diffused from its binding site inside UCP1 into the matrix leaflet in four out of six simulations (Fig. 2D).

For a complete proton transfer from the IMS to the matrix, the FAp must release its proton to the matrix. To explore the potential for FAp to lose its proton to the matrix (Fig. 2E), we launched constant pH simulations at the matrix pH(=7.8).

### Cationic UCP1 Residues Facilitate Fatty Acid Deprotonation

The initial configurations for the 1 µs constant pH simulations were derived from snapshots of previous simulations where FAp was either near R71 (simulation CpH-R71) or adjacent to TM6 (simulation CpH-TM6). All lysine, glutamate, and histidine residues facing the matrix (K56, K67, K73, K151, K175, K249, K257, K269, E46, E69, E168, E262, H146, H148) were kept titratable.

In the CpH-R71 simulations, FAp deprotonated to FA*^−^* within 100 ns and remained deprotonated for the remainder of the simulation (Fig. S10). FA*^−^* bound R71 and K67, with its C2 atom positioned next to V66 (Fig. 5C), and remained in this position for the rest of the simulation. Interestingly, the binding of FA*^−^* between K67 and R71 correlates well with an undetermined lipid density in the apo-UCP1 cryo-EM structure [Kang and Chen, 2023] (Fig. S11). The undetermined density in the cryo-EM structure probably represents a fatty acid or an anionic phospholipid.

In the CpH-TM6 simulation, FAp diffused to R71 after 550 ns and then got deprotonated. The resulting FA*^−^* remained bound to R71 for ∼100 ns before diffusing into the membrane (Fig. S10). After diffusing within the matrix leaflet for ∼150 ns, the FA*^−^* got protonated again, indicating that the presence of cationic residues on UCP1 is critical for inducing deprotonation of FAp. The pKa of FAp decreases in the presence of such cationic residues on UCP1, but increases again when the FA*^−^* diffuses into the membrane.

So far, our data has shown that (1) FA*^−^* can spontaneously approach D28*^−^* in the central UCP1 substrate-binding site, (2) becomes protonated via a water molecule bridging D28*^−^* and FA*^−^*, (3) diffuse as FAp to the matrix leaflet and then (4) deprotonates in the matrix leaflet.

The final step in the proton transfer cycle is the return of the deprotonated FA*^−^* from the matrix leaflet to the IMS leaflet (Fig. 2F). A spontaneous transmembrane flip is energetically unfavorable for the anionic FA*^−^*, with the barrier of a flip previously estimated to be ∼20 kcal/mol [Kampf et al., 2006]. We speculated that UCP1 could lower the free energy barrier by providing a polar pathway for FA*^−^* to diffuse back to the IMS. The reduction of the free energy barrier for the transmembrane transport of hydrophilic solutes due to the presence of a polar pathway on the membrane-protein interface of transmembrane proteins is not unprecedented. A similar mechanism was recently reported for a mitochondrial outer membrane protein insertase, which facilitates the transport of a lipid by lowering the energy barrier [Barto_̌_s et al., 2024]. We examine this possibility for UCP1 in the following sections.

### ATP Disrupts the Membrane-Facing Return Route for FA**^−^**

To identify a potential transport pathway for FA*^−^* to return to the IMS, we launched three 20 µs coarse-grained simulations of UCP1. In these simulations, UCP1 was embedded in an IMM containing 11 FA*^−^* molecules randomly distributed in the matrix leaflet. Coarse-grained simulations allow faster sampling of interactions between FA*^−^* and the UCP1 surface compared to all-atom simulations, and are often used to analyse lipid-protein interactions in transmembrane proteins [Corradi et al., 2019]. Although no FA*^−^* spontaneously flipped to the IMS, UCP1 facilitated FA*^−^* penetration ∼4 Å deeper into the membrane compared to simulations of an IMM without UCP1. FA*^−^* was frequently located along a polar stretch on TM2 (Fig. 6A,B), which accounted for the deeper membrane penetration of the FA*^−^* with UCP1. One of the residues on the polar stretch: Q83, exhibited the highest FA*^−^* occupancy, despite being positioned at the center of the membrane (Fig. 6A). Additionally, residues T134 and R162 in the polar pathway had increased FA*^−^* occupancies. Cationic R162 is likely to initially anchor FA*^−^* to the polar pathway.

**Figure 6:**
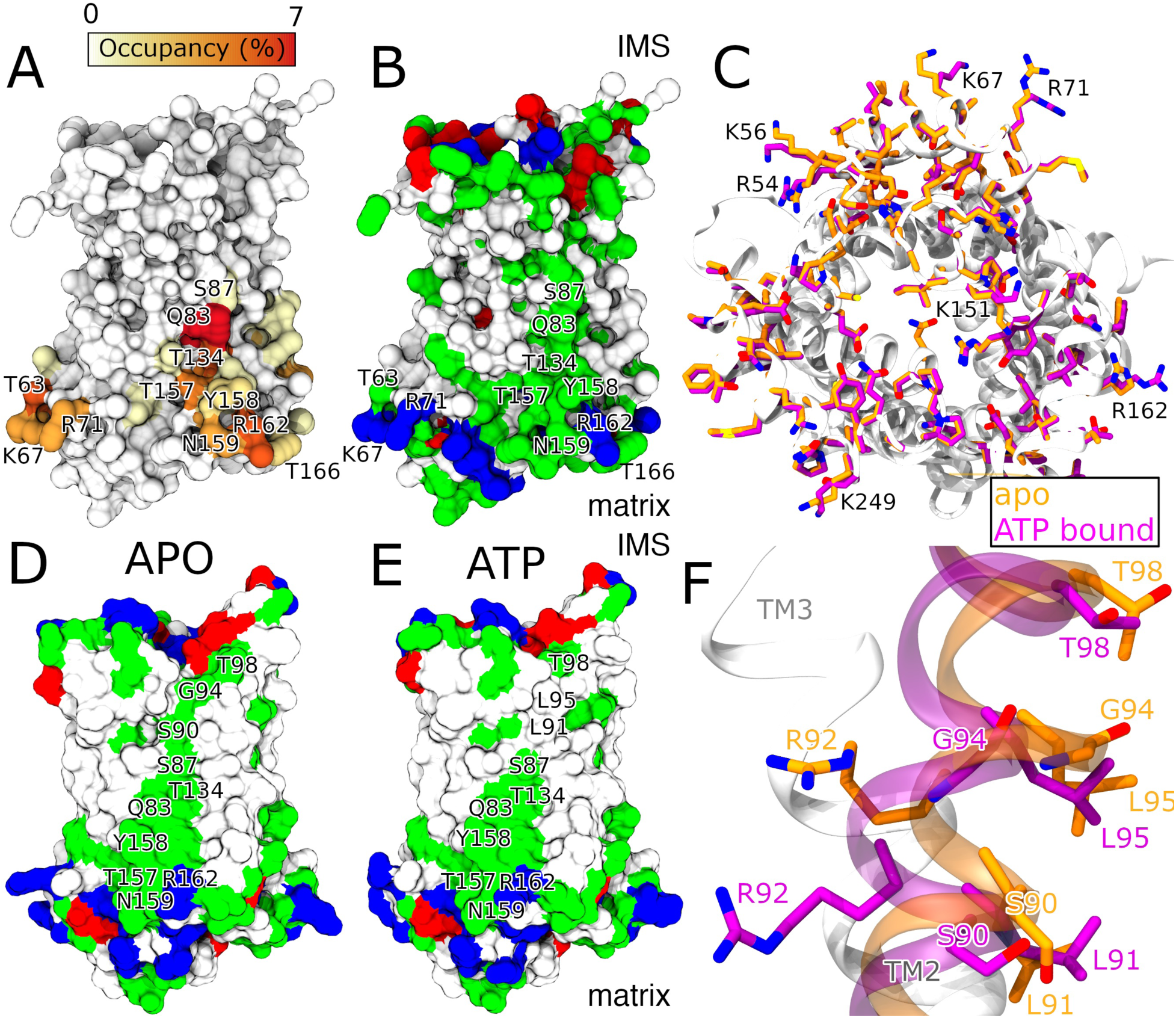
FA*^−^*-UCP1 interaction along a polar pathway. **A**: The interaction occupancy of FA*^−^* on the exterior of UCP1, averaged over the last 15 µs of three 20 µs coarse-grained simulations with 11 FA*^−^* molecules in the matrix leaflet. Red indicates ∼7% occupancy, while white indicates zero occupancy. **B**: Same UCP1 structure as in **A**, with residues colored by chemical properties: white (hydrophobic), green (polar), blue (basic), and red (anionic). **C**: Matrix-facing view of UCP1 in the cryo-EM structures of apo (orange; pdb 8hbv) and ATP-bound (purple; pdb 8hbw) states. **D,E**: Surface representation of UCP1 from the cryo-EM structures of the apo and ATP-bound states with residues colored by chemical properties as in **B**. **F**: TM2 in the cryo-EM structure of apo (orange) and ATP-bound (purple) UCP1.

We identified an identical polar pathway extending along TM2 from the matrix to the IMS in the cryo-EM structure of apo-UCP1 (pdb 8hbv, Fig. 6D). However, this polar pathway is interrupted between residues S87 and T98 in the ATP-bound (pdb 8hbw, Fig. 6E) and GTP-bound (pdb 8g8w) states [Kang and Chen, 2023, Jones et al., 2023]. We hypothesised that the continuous polar pathway in apo-UCP1 (absent in the nucleotide-bound states) could facilitate the return of FA*^−^* to the IMS leaflet. In the nucleotide-bound state, the nucleotide induces a local rotation of TM2 by binding to R92. This rotation replaces polar residues S90 and G94 on the UCP1 surface with hydrophobic residues L91 and L95 (Fig. 6D-F). Such a rotation is absent in the apo state [Jones et al., 2024, Kang and Chen, 2023].

Compared to all other helices, TM2 rotates the most upon binding of ATP. The RMSD of the TM2 backbone in the ATP-bound state relative to apo-UCP1 was 1.5 nm, while the corresponding RMSD values for the other five TMs were *<*0.55 nm. The higher RMSD of TM2 relative to apo-UCP1 was preserved in the CpH-APO and CpH-ATP simulations (Fig. S12). To probe this rotation further, we performed a principal component analysis (PCA) of the protein backbone (excluding loops and linkers).

PCA of the CpH-ATP, CpH-APO, and CpH-FA simulations showed that the ATP-bound UCP1 remained in a single conformational state in which the polar pathway was disrupted (Figs. S13 and S14). In contrast, 2-3 distinguishable conformational states were identified in both the FA-bound and apo states. Each of these conformations maintained the continuous polar pathway. Movement along eigenvector 1 in the apo state showed a local rotation of TM2 around R92 (Supplementary video 2). This rotation was not observed in the ATP-bound state, corroborating that TM2 loses flexibility upon nucleotide binding. Therefore, in both static cryo-EM structures and simulations, the apo and FA-bound states maintain a polar pathway along TM2 at the membrane-protein interface, while the pathway is disrupted in the nucleotide-bound state.

We hypothesized that the disruption of the polar pathway along TM2 in the presence of ATP has significant consequences on FA*^−^* transport from the matrix to the IMS. To quantify if UCP1 could indeed facilitate the flipping of FA*^−^*, we calculated the free energy barrier of FA*^−^* flipping from the matrix to the IMS under different conditions: with and without UCP1, and with or without ATP.

### UCP1 Facilitates FA**^−^** Transport from the Matrix to the IMS

We conducted umbrella sampling [Torrie and Valleau, 1974, 1977] simulations to generate potential of mean force (PMF) [Sprik and Ciccotti, 1998, Jain, 1997] profiles for the translocation of FA*^−^* across the membrane for three systems: apo-UCP1, ATP-bound UCP1, and a control system which is a membrane without UCP1.

FA*^−^* preferentially partitioned from the matrix into the matrix leaflet, indicated by a free energy minimum in all three systems (B in Fig. 7). At this position, FA*^−^* adopted an orientation similar to the membrane lipids, with the negatively charged carboxyl group facing the solvent and the hydrophobic tail pointing along the membrane lipid tails. However, further displacement of the charged carboxyl group into the hydrophobic membrane core was energetically unfavorable, as reflected by the rising PMF profiles beyond the free energy minimum toward the membrane center (B→C in Fig. 7).

**Figure 7:**
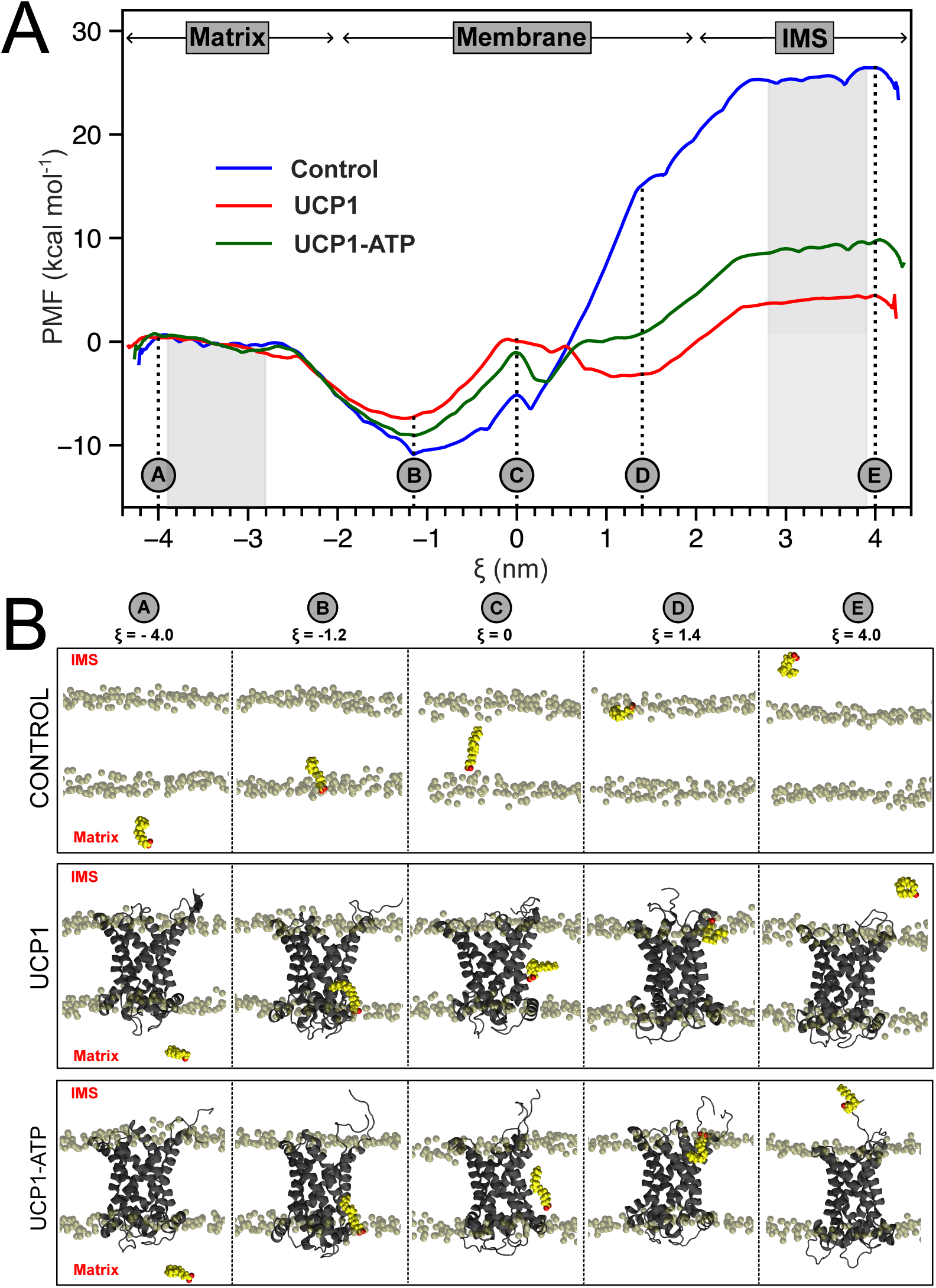
Potential of mean force (PMF) for FA. *^−^* **transport across the membrane. A**: PMF profiles for FA*^−^* transport from the matrix to the IMS with the distance of FA*^−^* from the membrane center as the reaction coordinate (*ξ*). The shaded regions were used to calculate mean PMF values, representing FA*^−^* in the matrix and IMS. **B**: Representative simulation snapshots corresponding to selected *ξ* values (A-E), illustrating key stages of FA*^−^* translocation.

The PMF profiles of FA*^−^* translocation into the matrix leaflet were similar across all systems, regardless of the presence of UCP1. However, significant differences emerged when FA*^−^* moved toward the IMS leaflet. In the control system, the PMF profile showed a sharp increase beyond the membrane center (C→D in Fig. 7). This increase resulted from the unfavorable orientation of the charged FA*^−^* within the hydrophobic membrane. As the FA*^−^* approached the lipid head groups in the IMS leaflet, it flipped such that the carboxyl group interacted with the membrane lipid headgroups. Subsequent movement of FA*^−^* from the IMS leaflet into the IMS resulted in a gradual rise in the PMF profile, eventually converging to a constant value in the solvent.

The presence of UCP1 reduced the energetic cost of the FA*^−^* flip significantly (Fig. 7). The FA*^−^* carboxyl group interacted with polar residues on TM2, reducing the energetic penalty of keeping a carboxyl group in the hydrophobic membrane. The PMF profile had a broad minimum around D in Fig. 7 for apo-UCP1. This minimum arises because polar residues S90 and G94 in the apo state (but not ATP-bound state; Fig. 6D,E) aid in the flipping and stabilization of FA*^−^* in the IMS leaflet.

The PMF profiles for the exit of FA*^−^* from the IMS leaflet into the solvent were similar across all three systems. The free energy barrier for FA*^−^* transport across the membrane was 25.5±0.7 kcal/mol without UCP1. This barrier was reduced to 4.3±1.2 kcal/mol in the presence of apo-UCP1, whereas ATP-bound UCP1 approximately doubled the barrier to 9.3±1.6 kcal/mol. Hence, the PMF data show that apo-UCP1 significantly reduces the free energy barrier of FA*^−^* transport, while the presence of ATP more than doubles the barrier compared to apo-UCP1.

### Mutational Analysis of Key Residues in UCP1

To validate the proton transport mechanism proposed by the simulations, we evaluated the proton conductance in UCP1 variants where key residues implicated in the simulations were mutated. Specifically, we studied the single mutations D28N, R71L, and Q83L, as well as the double mutant S90L/G94L. In Figure 8, green squares highlight the mutated residues across different UCPs and species. Residue D28 facilitates the protonation of FA*^−^* in the central substrate binding site (Fig. 2C), and R71 promotes the release of FAp’s proton to the matrix (Fig. 2E). Q83, S90, and G94 contribute to the polar pathway guiding FA*^−^* back to the IMS leaflet (Fig. 2F). Q83 showed the highest interaction occupancy with FA*^−^* (Fig. 6A), and S90 and G94 are replaced by leucine on the UCP1 surface in the ATP-bound state (Fig. 6D,E).

**Figure 8:**
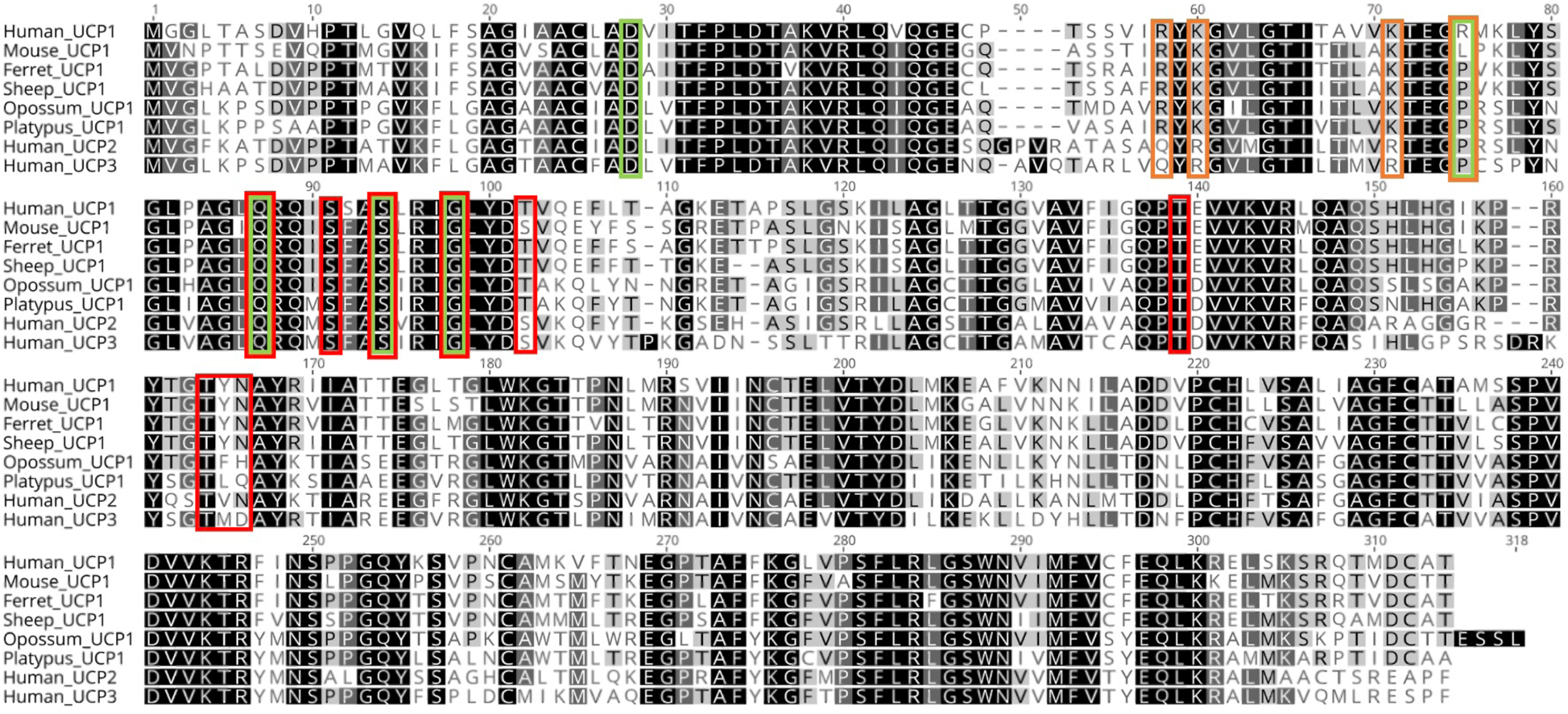
UCP1 amino acid alignment. Multiple sequence alignment of human UCP1 (Accession: NM 021833.5), UCP2 (Accession: U82819.1), UCP3 (Accession: NM 003356.4), and UCP1 from mouse (*Mus musculus*; Acces-sion NM 009463.3), ferret (*Mustela putorius furo*; Accession: AEYP01069989.1), sheep (*Ovis aires musimon*; Accession: CBYI010017988.1), opossum (*Monodelphis domestica*; Accession: OP589293.1), and platypus (*Ornithorhynchus anatinus*; Accession: AAPN01365328.1 and AAPN01365324.1). Alignment shading indicates regions of 100% similarity (black), 80-99% similarity (dark grey), 60-79% similarity (light grey), and less than 60% similarity (white). Squares highlight residues targeted in the mutation study (green), residues in the polar pathway (red), and basic residues on the repeat connecting TM1 and TM2 (orange).

We transiently overexpressed human UCP1 (hUCP1) mutants in human embryonic kid-ney (HEK293) cells. Immunoblotting revealed poor expression of the D28N and S90L/G94L mutants, suggesting detrimental effects on protein stability or increased degradation (Fig. S15). In contrast, the R71L and Q83L mutants displayed expression levels similar to wild-type (wt) hUCP1 (Fig. 9A,B). Only mutants with expression levels similar to wt hUCP1 were func-tionally tested (Fig. 9C-H).

**Figure 9:**
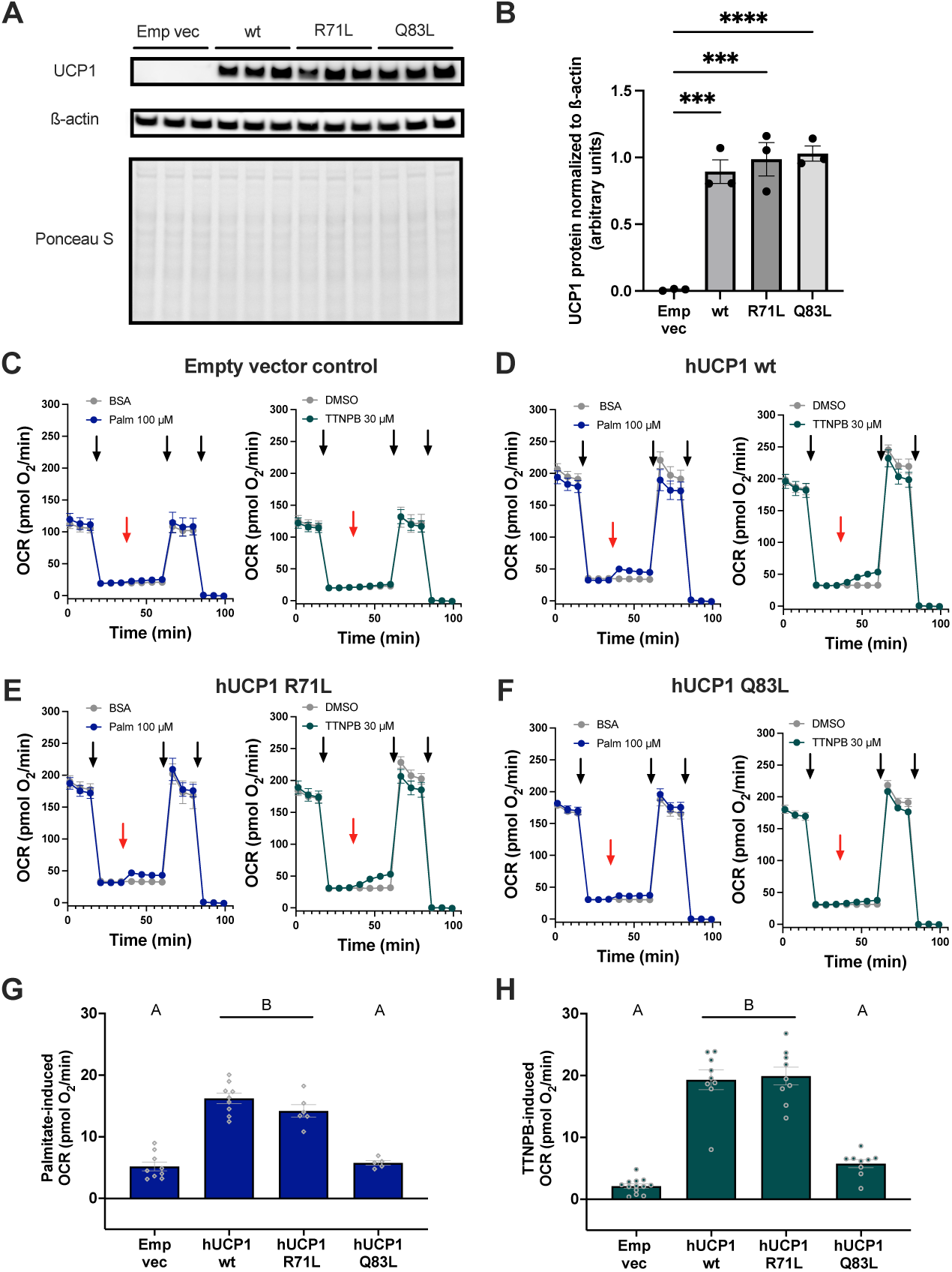
UCP1 protein quantification and plate-based respirometry. **A**: Western blot of protein lysates from HEK293 cells transfected with an empty vector (Emp vec) or wild-type (wt) and mutant (R71L or Q83L) human UCP1 (hUCP1). **B**: Densitometric UCP1 protein quantification across transfections, normalized to *β*-actin (***P *<* 0.001, ****P *<* 0.0001; n = 3 independent transfections for each variant). Averaged plate-based respirometry traces of intact HEK293 cells transfected with **C**: an empty vector control, **D**: wt UCP1, **E**: the R71L mutant, and **F**: the Q83L mutant. Injections of oligomycin, DNP, and rotenone/antimycin A during respirometry are indicated from left to right by black arrows, respectively. Red arrows signify the injections of vehicle controls (BSA or DMSO) or UCP1 activators (palmitate [palm]) or TTNPB). n=5-11 measurements performed on three independent days. Bar charts quantifying the oxygen consumption rates (OCR) response to **G**: palmitate and **H**: TTNPB. Statistically different groups (P *<* 0.0001) are denoted by letters A and B. Data are presented as means±SEM.

Plate-based respirometry was employed to measure cellular oxygen consumption rates (OCRs) of cells with wt and mutant hUCP1 as well as a control system [Jastroch et al., 2012, Gaudry and Jastroch, 2021]. Throughout the runs, discrete modules of respiration were delineated by the sequential injection of chemical activators and inhibitors [Divakaruni et al., 2014]. Following the measurement of basal respiration, oligomycin was injected to inhibit ATP synthase and thereby ATP-linked respiration. The residual respiration reflects a combination of proton leak and non-mitochondrial respiration. UCP1 variants were subsequently activated by free fatty acids (palmitate) or the non-canonical activator (arotinoid acid [TTNPB], a retinoic acid analog) [Tomás et al., 2004], resulting in increased OCR due to proton leak. Maximal respiration was then induced with an artificial uncoupler (e.g. 2,4-dinitrophenol). Finally, mitochondrial respiration was inhibited using rotenone and antimycin A to inhibit complexes I and III of the electron transport chain, respectively, enabling correction for non-mitochondrial respiration.

The plate-based respirometry revealed that wt hUCP1 significantly increased the OCR in response to both TTNPB and palmitate, compared to control cells (Fig. 9C-D,G,H). The R71L mutant exhibited an OCR response to palmitate and TTNPB similar to wt hUCP1, suggesting that R71L does not impair palmitate-or TTNPB-induced proton leak (Fig. 9D,E,G,H). In contrast, the Q83L mutation abolished the palmitate- and TTNPB-induced OCR (Fig. 9F,G,H), making it indistinguishable from the control system (Fig. 9C,G,H).

## Discussion

### The Binding Conformation of FA**^−^** in the UCP1 Central Binding Site

Starting from the cryo-EM structures of the apo and ATP-bound states of UCP1, we present a new model for the transport of protons and FA in UCP1 (Fig. 2). All-atom simulations revealed that FA*^−^* spontaneously diffused to and bound the central substrate-binding site of UCP1. Once bound, the FA*^−^*’s carbon chain extended into the IMM through the gap between TM5 and TM6 (Fig. 3D). Bertholet et al. reported a similar FA*^−^* binding conformation in seven 2-5 µs simulations of FA*^−^* bound to ANT, another member of the SLC25 family [Bertholet et al., 2022]. In their simulations, FA*^−^* interacted with residues K22, R79, and R279, which correspond to D28, R84, and R277 in UCP1, respectively. The FA*^−^* tail also extended into the membrane from between TM5 and TM6 in their simulations. Hence, FA*^−^* binds in a similar manner in UCP1 and ANT.

Prior binding kinetics of mant-GDP and enzymatic proteolysis indicated that UCP1 changes conformation upon FA-binding with some suggesting that the conformational change was from a c-state to an m-state [Divakaruni et al., 2012]. Specifically, proteolysis targeting K292 on TM6 was accelerated upon FA-binding. Based on our FA-bound UCP1 conformation, we argue that the accelerated proteolysis can be due to a small conformational change of TM6, as the FA tail extends into the IMM between TM5 and TM6, making K292 more accessible to the degrading enzyme during proteolysis. Additionally, binding kinetics was unchanged for other nucleotide analogues (mant-ATP, mant-ADP or mant-GTP) suggesting that the conformational change must be subtle, to only influence GDP-binding kinetics. The c-state to m-state conformational change should be more drastic and significantly alter the binding kinetics of all nucleotides. Protein stability analysis indicates that UCP1 undergoes a destabilizing conformational change upon FA-binding [Cavalieri et al., 2022], but there is no indication that the m-state is involved. We did not observe any significant conforma-tional changes upon FA-binding in our simulations, which may not be long enough to sample conformational changes which occur on longer timescales.

### Role of D28 and Water Molecule Coordination in Protonation of FA**^−^**

When negatively charged FA*^−^* or ATP binds to UCP1, the electrostatic environment of the central substrate-binding site is altered, increasing the likelihood of D28 attracting a proton. Initially, we hypothesized that protonated D28p would donate a proton directly to FA*^−^*. However, in our all-atom simulations, FA*^−^* never approached D28p close enough to effect such a transfer. Instead, we found that a water molecule frequently bridged FA*^−^* and D28*^−^*. We suspected that the water molecule could potentially effect the proton transfer.

Protonation of FA*^−^* via the water molecule was directly observed in QM/MM simulations, and FEP calculations showed that the proton transfer from D28p to FA*^−^* was energetically favorable with a free energy difference of −1.3 ± 0.4 kcal/mol. These results support our postulate that the close proximity of the D28*^−^* and FA*^−^* carboxylate groups creates a negatively charged environment that attracts a proton from the IMS which protonates either D28*^−^* or FA*^−^*. Kunji and Robinson made a similar hypothesis for anionic substrates and mitochondrial carriers in general [Kunji and Robinson, 2010]. Their hypothesis was based on the observation that mitochondrial carriers facilitating substrate-proton symport contain an anionic residue at the substrate-binding site [Kunji and Robinson, 2010]. The hypothesis is supported by the surplus of protons in the IMS, which can be readily attracted to the negatively charged site between the carboxylate groups.

The direct involvement of D28 in proton transport has been documented several times [Klingenberg et al., 1999, Echtay et al., 2000, Urbánková et al., 2003a]. Mutations D28N and D28V reduced proton transport to 8-60% of wt levels. In contrast, proton transport was unaffected by the D28E mutation, which underlines the critical role of the carboxylate anion [Echtay et al., 2000]. D28 is proposed to be functionally equivalent to D134 in the substrate-binding site of ANT. Indeed, the D134S mutant in ANT reduced proton transport by 60% [Kreiter et al., 2023].

### Role of the R277-D28 Salt Bridge in FA**^−^** Binding and Proton Transfer in UCP1

Our constant pH and standard all-atom simulations demonstrated that FA*^−^* only approached D28*^−^* when a salt bridge was formed between R277 and D28*^−^*. When FA*^−^* approached D28*^−^*, R277 simultaneously bound FA*^−^* and D28*^−^* (Figs. S4 and S7). Thus, R277 plays a dual supporting role in the proton transfer: it is a proton surrogate for D28*^−^*, as we proposed previously [Jacobsen et al., 2023], and helps FA*^−^* attraction towards D28*^−^*. The D28-R277 salt bridge is present in all currently resolved ligand-bound UCP1 cryo-EM structures (ATP [pdb 8hbw], GTP [pdb 8g82], and DNP [pdb 8j1n]) but is absent in the apo state (pdb 8hbv), as was also recently noted by Jones et al. [Kang and Chen, 2023, Jones et al., 2023, 2025]. This observation suggests that R277 binds D28 only in the presence of a negatively charged ligand. The observations we have made about R277 are in conflict with the finding that the charge-neutralizing R277Q mutation does not impair proton transport [Echtay et al., 2001]. We speculate that another cationic residue from the arginine triplet, such as R84 fulfils the role of R277 in the R277Q mutation.

### Role of Basic Residues in FA**^−^** Deprotonation

After protonation, FAp exited UCP1 between TM5 and TM6. FAp tended to interact with R71 in both all-atom and coarse-grained simulations, and the cationic residue induced a pKa shift which lead to deprotonation of FAp.

On the matrix facing side of UCP1, six basic residues extend into the membrane (R54, K56, K67, R71, R162, and K249) of which four (R54, K56, K67, and R71) are concentrated on the repeat connecting TM1 and TM2 (Figs. 6C). The repeat connecting TM1 and TM2 aligns with the exit route of FAp (between TM5 and TM6; Fig. 1B). These basic residues can facilitate FAp deprotonation, and the resulting FA*^−^* can bind them, e.g., between R71 and K67 as observed in our simulations (Fig. 5C). Along these lines, the mutations K48S, R59S, and K62S in ANT reduced proton transport through ANT by ∼70%. Similarly, simulations of ANT indicated tight interactions between FA*^−^* and R59 and K62 (K67 in UCP1) [Kreiter et al., 2023].

Like R54, K56, K67, and R71 in UCP1, K48, R59, and K62 in ANT are located on the repeat connecting TM1 and TM2. We predict that mutations of R54, K56, K67, and R71 in UCP1 should reduce proton transport. However, the OCR assays showed that fatty acid-induced proton transport was unaffected by the R71L mutation (Fig. 9D,E). R71 is not conserved across UCP1 species and is often replaced by proline (Fig. 8). This suggests that R71 alone is unlikely to mediate FAp deprotonation in the matrix leaflet. Instead, the combined effect of multiple basic residues (R54, K56, K67, and R71) at the same site promotes the deprotonation. Unlike R71, the residues R54, K56, and K67 are conserved across UCP1 species (Fig. 8), highlighting their potential functional significance. Further corroboration of our finding that FA*^−^* binds to R71 and K67 in hUCP1 comes from the unassigned lipid density along TM1 in the apo-UCP1 cryo-EM structure (Fig. S11).

### Nucleotide Binding Inhibits FA**^−^** Transport by Disrupting a Polar Pathway along TM2

Umbrella sampling calculations showed that the free energy barrier for the return of FA*^−^* from the matrix to the IMS leaflet is significantly reduced in the presence of UCP1. Apo-UCP1 provides a polar pathway along which FA*^−^* can slide back to the IMS. However, this polar pathway is disrupted by nucleotide binding, revealing a novel mechanism by which nucleotides can inhibit proton transport in UCP1. Although the barrier is reduced to 4.3 kcal/mol in apo-UCP1, FA*^−^* will still not flip spontaneously to the IMS even in the presence of UCP1. However, the magnitude of the barrier can be significantly reduced by the transmembrane electrical gradient across the IMM which will aid the transport of FA*^−^* from the matrix to the IMS.

Residues Q83, S87, S90, G94, T134, and T157 within the polar pathway are conserved across hUCP1-3 and various UCP1 orthologs (Fig 8). The mutation Q83L abolished FA-induced proton leak by UCP1 (Fig. 9), demonstrating that the polar pathway and specifically interaction with Q83 is vital for FA-induced proton transport. T98 is sometimes substituted by a serine, which will preserve the polar pathway between Q83 and T98. A polar pathway along TM2 is, therefore, also visible in the AlphaFold-predicted structures of human UCP2 (https://alphafold.ebi.ac.uk/entry/P55851) and UCP3 (https://alphafold.ebi.ac.uk/entry/P55916) [Jumper et al., 2021, Varadi et al., 2022]. One of the first residues of the polar pathway, Y158, is substituted by hydrophobic amino acids (valine, methionine, leucine, or phenylalanine) in human UCP2 and UCP3, and in non-thermogenic UCP1 orthologs such as those from platypus and opossum (Fig 8). This substitution can explain the significantly lower proton transport rates in UCP2 (4.5 protons/s) [Beck et al., 2007] and UCP3 (2.6 protons/s) [Macher et al., 2018] compared to thermogenic UCP1 (14 protons/s) [Urbánková et al., 2003b]. In these proteins, the polar pathway appears disrupted at its onset, thus compromising proton transport rates.

Additionally, N159 is conserved in thermogenic UCP1. In non-thermogenic UCP1 and in human UCP2 and UCP3, N159 is typically replaced by a different polar residue and, therefore, the effect of such a substitution on FA*^−^* transport is unclear. Intriguingly, in human UCP3, N159 is replaced by anionic D159 (Fig. 8). Positioned adjacent to R162, D159 is likely to form a salt bridge with R162, resulting in charge screening. This interaction will interfere with the initial anchoring of FA*^−^* to the polar pathway, and can explain the reduced proton transport capacity of UCP3 compared to UCP2 and, particularly, UCP1. In conclusion, we have put forward a novel mechanism for proton and FA transport through UCP1, which differs from the previously proposed Buffering, Shuttling, and Cycling models. Our mechanism explains every step of the transport cycle, is supported by simulation and biochemical data, and explains a diverse set of biochemical data about the transport mechanisms in UCP1 and its analogues: ANT, UCP2 and UCP3.

**Table 1:**
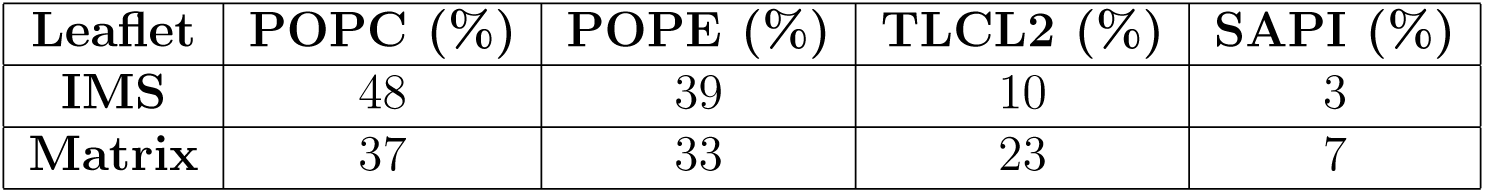
Membrane composition used in the simulations to mimic the IMM. POPC: 3-palmitoyl-2-oleoyl-glycero-1-phosphatidylcholine, POPE: 3-palmitoyl-2-oleoyl-glycero-1-phosphatidylethanolamine, TLCL2: tetralinoleoyl cardiolipin (head group charge=-2), SAPI: 3-stearoyl-2-arachidonoyl-glycero-1-phosphoinositol.

## Methods

In all simulation setups, three cardiolipin lipids were pre-positioned at their known binding sites on UCP1, and the membrane composition was designed to mimic an IMM (Table 1) [Daum and Vance, 1997, Horvath and Daum, 2013]. Palmitoleic acid (C16:1) was used as the representative FA. All simulations, except for QM/MM, were performed with Gromacs 2021.1 [Abraham et al., 2015, 2024] or newer versions. The CHARMM36m force field[Huang et al., 2017, Brooks et al., 2009b, Klauda et al., 2010] was used. Modeller [Sali and Blundell, 1993] was used to add missing termini in the experimental structures (residues M1-V9 and R300-T307). FA-UCP1 interactions were quantified using ProLIF [Bouysset and Fiorucci, 2021] and PyLipID [Song et al., 2021]. PCA clustering was performed with the OPTICS method in the Scikit-learn Python library[Pedregosa et al., 2011], using a minimum neighborhood size of 2000 samples and a minimum cluster boundary steepness of 0.001. Molecular graphics were generated using VMD [Humphrey et al., 1996].

### All-Atom Molecular Dynamics

All-atom simulations were initially prepared using the membrane builder function in CHARMM-GUI [Jo et al., 2008, Brooks et al., 2009a,b, Jo et al., 2007], while minor adjustments, such as adding FAs or neutralizing ions, or altering protonation states were performed manually. The simulation systems included ∼23,000 water molecules and a 0.15 M NaCl concentration. Systems were minimized using 5,000 steps of the steepest descent algorithm and equilibrated over five steps, during which restraints on the system were gradually relaxed. Exceptions were made for simulations that continued from already equilibrated systems with minor modifications. Production simulations were run for 500-5000 ns with a 2 fs time step at 1 atm, using the semi-isotropic Parrinello-Rahman barostat [Parrinello and Rahman, 1981] with a time constant of 5 ps and a compressibility of 0.00045/bar. The temperature was maintained at 310 K using the Nośe-Hoover thermostat [Nośe, 1984, Hoover, 1985] with a time constant of 1 ps. The Linear Constraint Solver [Hess et al., 1997] algorithm was applied to constrain all bonds involving hydrogen atoms. A 1.2 nm cutoff was used for short-range interactions, and particle mesh Ewald [Darden et al., 1993, Essmann et al., 1995] was employed for long-range electrostatic interactions.

### Constant pH Simulations

Constant pH simulations were conducted using the Gromacs constant pH code developed by Aho et al. [Aho et al., 2022, Buslaev et al., 2022]. These simulations employed the same MD settings as the standard all-atom simulations. The pH was set to 6.8 or 7.8, depending on whether residues facing the IMS or the matrix were studied, respectively. To accommodate the changes in charge associated with protonation shifts in the protein or FA, 40 buffer particles with a mass of 5 were added to the solvent. These buffer particles had and initial charge of 0 but could carry charges ranging from −0.5 to +0.5.

For amino acids, the reference pKa values and the coefficients of the fit to *∂V* ^MM^*/∂λ* (where *V* ^MM^ is a correction potential) were adopted from the original paper by Aho et al. [Aho et al., 2022, Buslaev et al., 2022]. The FA reference pKa was 5, and the *∂V* ^MM^*/∂λ* coefficients were determined following the parameterization protocol described in the original publication [Aho et al., 2022] and its associated manual [Aho et al., 2021]. The barrier of the biasing potential, which enhances sampling of the physical states at *λ* = 0 and *λ* = 1, was set to 6 kJ/mol for amino acids and the FA, and 0 kJ/mol for the buffer particles.

CpH-APO and CpH-ATP simulations were based on the 300 ns PRE-APO and PRE-ATP simulations, respectively, while the CpH-FA simulation continued from a 1100 ns simulation of an FA*^−^* bound UCP1. For titration curve simulations (Fig. S3), constant pH simulations of 100 ns were performed at each pH value from 1 to 9, in increments of 1.

### Quantum Mechanical/Molecular Mechanics Simulations

QM/MM simulations were performed using Amber22 [Case et al., 2022]/Terachem [Seritan et al., 2021] (version 1.96H), with the QM region treated at a CAM-B3LYP [Yanai et al., 2004]-D3(BJ) [Grimme et al., 2010, 2011]/def2-SVP [Weigend and Ahlrichs, 2005] level of theory. MM parameters were transferred from the previous classical MD simulations and converted from Gromacs to Amber format using Parmed. The QM region consisted of the carboxyl group and the three next carbons along the chain of the fatty acid (and connected hydrogens), the D28 side chain, and a single water molecule. Constant-volume QM/MM simulations were run for 3-45 ps, using a 1 fs time-step. The temperature was controlled towards 310 K using a Langevin thermostat.

### Free Energy Perturbation

The FEP simulations employed the same MD settings as the standard all-atom simulations. Harmonic distance restraints with a force constant of 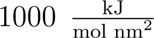 were applied between the oxygen atom of the coordinated water molecule and the oxygen atom of D28 (2.97 Å), as well as between the water molecule and the FA (2.65 Å). The difference in free energy due to the restraints was negligible, as detailed in the supplementary material (’Free Energy Perturbation’). The system was first minimized using 5000 steps and then equilibrated for 2 ns.

The proton transfer was performed with 20 equidistant *λ*-windows (interval size 0.05), with each window consisting of a 20 ns simulation. Restraint decoupling was performed dur-ing 48 *λ*-windows, with the first 40 having a window size of 0.025 and the last 10 windows having a reduced interval size of 0.005 to allow for improved sampling near the complete decoupling of the restraints. For restraints each window was simulated for 25 ns. Restraints were decoupled linearly with *λ* while the soft-core potential introduced by Beutler et al. [Beutler et al., 1994] was applied during the the proton transfer FEP with the soft core parameter set to 0.7 and the *λ* power set to 1. The final 10 ns of each window were used for analysis. The free energy measure was based on the forward and backward transformation data and was calculated using the Bennet Acceptance Ratio [Bennett, 1976].

In the final *λ*-window of the restraint decoupling, a flat-bottom potential was applied to address periodic boundary condition issues caused by free diffusion of the water molecule. This potential had a reference distance of 3 nm and a force constant of 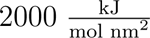 between the *α*-carbon of D28 and the coordinated water molecule.

The thermodynamic cycle used for FEP and the free energy change as a function of *λ*-windows are shown in Figs. S16 and S17. There is sufficient sampling within each *λ*-window (Figs S18 and S19) and good phase space overlap between neighboring *λ*-windows (Figs. S20-S23).

### Coarse-Grained Molecular Dynamics

Coarse-grained simulations were prepared using the MARTINI bilayer maker in CHARMM-GUI [Jo et al., 2008, Brooks et al., 2009a,b, Qi et al., 2015]. Since parameters for cardiolipin and FA were unavailable in MARTINI3 [Souza et al., 2021], the simulations utilized MARTINI2 [Marrink et al., 2004, 2007] along with an elastic network model to restrain the tertiary structure of the protein. The solvent consisted of polarizable water with a 0.15 M NaCl concentration. The systems were minimized by 5,000 steps of the steepest descent algorithm and equilibrated during five stages during which restraints on the system were gradually relaxed.

Production simulations employed a 15 fs time step. The pressure was maintained at 1 atm using the semi-isotropic Parrinello-Rahman barostat [Parrinello and Rahman, 1981] with a time constant of 12 ps and a compressibility of 0.0003/bar. The system temperature was maintained at 310 K using the velocity rescale thermostat [Bussi et al., 2007] with a time constant of 1 ps. VdW and electrostatic interactions were cut off at 1.1 nm, and the short-range neighbor list was updated every 20 step. In these simulations, cardiolipin lipids were allowed to spontaneously diffuse and bind to UCP1.

### Umbrella Sampling Simulations

Umbrella sampling simulations were conducted to calculate the PMF for FA*^−^* translocation from the matrix to the IMS across three systems: the control system (membrane-only), apo-UCP1, and ATP-bound UCP1. In the control system, conformations for umbrella sampling were prepared using two separate pulling simulations: starting from the same initial position in the matrix leaflet, FA*^−^* was first pulled along the membrane normal into the solvent on the IMS side, and second, into the solvent on the matrix side. A force constant of 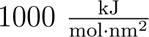 and a pulling rate of 0.0001 nm/ps were used.

Three 500 ns all-atom simulations of UCP1 embedded in an IMM with 12 FA*^−^* in the matrix leaflet revealed that residues R162 and R71 were the main interaction sites for FA*^−^*, with average occupancies of ∼50%, consistent with our previously published results [Jacobsen et al., 2023]. Based on this, pulling simulations in the apo-UCP1 and ATP-bound UCP1 systems began from conformations where FA*^−^* was bound to R162. To maintain proximity between FA*^−^* and the protein during pulling, a flat-bottom harmonic potential (force constant of 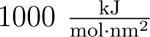) was applied in the membrane-plane between the center of mass (COM) of FA*^−^* and UCP1. For COM distances ≤2 nm, no potential was applied, while for distances *>*2 nm, the harmonic potential became active.

The initial conformations for umbrella sampling were extracted from the pulling trajectories with a spacing of 0.1 nm in the range ±4 nm from the membrane center, which ultimately produced ∼80 windows for each system. Each window was equilibration for 500 ps, followed by 50 ns production simulations. Umbrella sampling simulations employed the same MD settings as the standard all-atom simulations.

The potential of mean force was reconstructed using the weighted histogram analysis method [Kumar et al., 1992] and errors were estimated from 250 bootstrapped replicas. To calculate the free energy difference for FA*^−^* transport from the matrix to the IMS, the mean PMF value was averaged over the range 2.8-3.8 nm from the membrane center across all bootstrapped PMF profiles for the three systems.

### UCP1-Containing Vectors

Transient transfections of HEK293 cells were performed as described by Gaudry et al. [Gaudry et al., 2024]. Vectors containing hUCP1 variants (wt, single mutants D28N, R71L, Q83L, and the double mutant S90L/G94L) were synthesized by Azenta Life Sciences. The UCP1 coding sequences were cloned into the pcDNA3.1+ vector using Xhol and Xbal restriction sites. An empty pcDNA3.1+ vector was also prepared as a negative control. The vectors were first transformed in DH5*α E. coli* (Invitrogen), as per the manufacturer’s instructions, and grown overnight at 37*^◦^*C on LB agar plates with 100 µg/ml ampicillin. Single colonies were selected to inoculate liquid LB cultures that were grown at 37*^◦^*C overnight while shak-ing at 225 RPM. The plasmids were purified using a QIAprep Spin Miniprep Kit (Qiagen) and the sequence identities of the plasmids were verified by Sanger sequencing.

### HEK293 Cell Culture and Transfections

HEK293 cells were cultured at 37*^◦^*C and 5% CO_2_ in Dulbecco’s Modified Eagle Medium (DMEM), high glucose (Gibco) with 10% fetal bovine serum and penicillin/streptomycin (Gibco; 100 U/ml). To prepare the cells for transfection, they were first trypsinized from T175 flasks using Trypsin-EDTA (0.05%), phenol red (Gibco). The cells were then counted and diluted to a density of 1.76×10^6^ cells in 3.2 ml of growth medium. The transfection cocktails were prepared in a separate tube with 900 ml of pure DMEM, 36 µl PolyFect Transfection Reagent (Qiagen), and 9 µg of plasmid DNA. As specified in the manufacturer’s instructions, the transfection cocktails were mixed and then incubated at room temperature for 15 minutes before being added to the cell aliquots. The HEK293 cells were then seeded at a density of 30.000 cells per well on xf96 well plates that had previously been coated with polyethylenimine for plate-based respiratory experiments. Cells were also seeded in parallel on 12-well plates to verify protein levels using Western blots.

### Western Blotting

Cells seeded on 12-well plates were washed 3 times with PBS and frozen at −80*^◦^*C while plate-based respiratory experiments were taking place. Cell lysates were later collected on ice by adding 150 µl of RIPA buffer (50 mM NaCl, 50 mM TRIS, 0.5% sodium deoxycholate, 1% IGEPAL CA-630, and 0.1% sodium dodecyl sulfate) with HALT Protease and Phosphatase Inhibitor Cocktail (Thermo Scientific). The cell debris was pelleted by centrifuging the lysates at 18000×g for 30 minutes at 4*^◦^*C, and the supernatant was collected. The protein levels were quantified using Bradford reagent (Sigma-Aldrich) and bovine serum albumin protein standards. Wells of Bolt 4-12% Bis-Tris Plus gels (Invitrogen) were loaded with 10 µg of protein and electrophoresed in a Mini Gel Tank (Invitrogen) as described in the manufacturer’s protocol. Protein transfers to nitrocellulose membranes were performed using iBlot 2 NC Regular Stacks (Invitrogen) and the iBlot 2 Gel Transfer Device (Invitrogen). The membranes were rinsed in deionized water, then stained with Ponceau S to visualize total protein loading. The membranes were then blocked in 5% skimmed milk solution in Tris-buffered saline with Tween 20 (TBS-T) for 1 hour on a shaking platform at room temperature. The membranes were washed with TBS-T and the monoclonal anti-UCP1 antibody (MAB7158; R&D Systems) was applied overnight at 4*^◦^*C on a rocking platform. This antibody had been diluted 1:1000 in TBS-T with 5% bovine serum albumin (BSA; Sigma-Aldrich). The membranes were then washed with TBS-T and a 1:20000 dilution of goat anti-mouse IgG, HRP (AP130P; EMD Millipore Corp.) in 5% milk + TBS-T was added. After incubating at room temperature for 1 hour on a rocking platform, the membranes were imaged using Clarity Western ECL Substrate (Bio-Rad) on a Bio-Rad ChemiDoc MP. The blots were then stripped at room temperature for 20 minutes, blocked in 5% milk solution, and a 1:10000 dilution of *β*-actin HRP-linked antibody (Santa Cruz sc47778) in 5% milk + TBS-T was applied for 20 minutes before re-imaging the blot. Western blot densitometric analyses were performed using ImageLab 6.1 (Bio-Rad).

### Plate-Based Respirometry

Plate-based respirometry for intact HEK293 cells was performed as described by Gaudry et al. [Gaudry et al., 2024] (see reference for details) using a Seahorse XFe96 Extracellular Flux Analyzer (Agilent). Basal respiration was measured, followed by injection of oligomycin (end concentration: 4 µg/mL) to inhibit ATP-synthase-linked respiration. UCP1 mutants were activated using palmitate (end concentration: 100 µM) or the retinoic acid analog, arotinoid acid (TTNPB; end concentration: 3 µM). Injected vehicle controls for palmitate and TTNPB were fatty acid-free bovine serum albumin (BSA) or DMSO, respectively. Palmitate used for the experiments was equilibrated with BSA at a 6:1 molar ratio. Maximal respiration was verified with the injection of 2,4-dinitrophenol (DNP; end concentration: 100 µM). Lastly, a mixture of rotenone (end concentration: 4 µM) and antimycin A (end concentration: 2 µM) was injected to inhibit complexes I and III of the electron transport chain and correct for non-mitochondrial respiration. Traces were excluded if an injection port leaked prematurely into the experimental well.

All OCR were corrected for non-mitochondrial respiration. Bar charts of activator-induced OCR (Fig 9G,H) were calculated first by averaging the initial two measurement points after injection of palmitate, and the third and fourth measurement points after the injection of TTNPB. The average OCR of the three measurements following oligomycin injection were then subtracted from these palmitate- and TTNPB-induced OCR.

### Multiple Sequence Alignment

For the multiple sequence alignment, UCP1 nucleotide coding sequences were sourced from the NCBI RNA sequencing database or whole genome shotgun contigs. Accession numbers are listed in the caption of Fig. 8. The coding sequences were virtually translated to amino acid sequences in Geneious 9 (https://www.geneious.com). The alignment was performed with Multiple Sequence Comparison by Log-Expectation (MUSCLE [Edgar, 2004]).

### Statistics

Statistical analyses were performed in GraphPad Prism 10.1.1 for macOS (www.graphpad. com). Data are reported as means±SEM. One-way ANOVA was employed to assess differences between mutants with Tukey post hoc tests.

## Data Availability

TO BE MADE

## Acknowledgments

LJ and HK are supported by the Lundbeck foundation Ascending Investigator (grant R344-2020-1023). SM is supported by Danmarks Frie Forskningsfond | Natur og Univers (Grant 3103-00079B). AAHZ is supported by the Novo Nordisk Foundation (Grant NNF20OC0065368). MJG and MJ are supported by Vetenskapsrådet (Grants 2024-06689 and 2022-03136) and the Novo Nordisk Foundation (Grant 0059646). The simulations were performed on the Finnish Supercomputer LUMI, under grant numbers DeiC-AU-N5-2023014, DeiC-SDU-S5-202400009, and DeiC-SDU-S5-202400009, and on the Novo Nordisk Foundation-funded ROBUST Resource for Biomolecular Simulations (Grant NNF18OC0032608).

